# Reduced binding and neutralization of infection- and vaccine-induced antibodies to the B.1.351 (South African) SARS-CoV-2 variant

**DOI:** 10.1101/2021.02.20.432046

**Authors:** Venkata Viswanadh Edara, Carson Norwood, Katharine Floyd, Lilin Lai, Meredith E. Davis-Gardner, William H. Hudson, Grace Mantus, Lindsay E. Nyhoff, Max W. Adelman, Rebecca Fineman, Shivan Patel, Rebecca Byram, Dumingu Nipuni Gomes, Garett Michael, Hayatu Abdullahi, Nour Beydoun, Bernadine Panganiban, Nina McNair, Kieffer Hellmeister, Jamila Pitts, Joy Winters, Jennifer Kleinhenz, Jacob Usher, James B. O’Keefe, Anne Piantadosi, Jesse J. Waggoner, Ahmed Babiker, David S. Stephens, Evan J. Anderson, Srilatha Edupuganti, Nadine Rouphael, Rafi Ahmed, Jens Wrammert, Mehul S. Suthar

**Author notes:** Correspondence: Mehul S. Suthar and Jens Wrammert. Lead contact: Mehul S. Suthar.

## Abstract

The emergence of SARS-CoV-2 variants with mutations in the spike protein is raising concerns about the efficacy of infection- or vaccine-induced antibodies to neutralize these variants. We compared antibody binding and live virus neutralization of sera from naturally infected and spike mRNA vaccinated individuals against a circulating SARS-CoV-2 B.1 variant and the emerging B.1.351 variant. In acutely-infected (5-19 days post-symptom onset), convalescent COVID-19 individuals (through 8 months post-symptom onset) and mRNA-1273 vaccinated individuals (day 14 post-second dose), we observed an average 4.3-fold reduction in antibody titers to the B.1.351-derived receptor binding domain of the spike protein and an average 3.5-fold reduction in neutralizing antibody titers to the SARS-CoV-2 B.1.351 variant as compared to the B.1 variant (spike D614G). However, most acute and convalescent sera from infected and all vaccinated individuals neutralize the SARS-CoV-2 B.1.351 variant, suggesting that protective immunity is retained against COVID-19.

---

SARS-CoV-2 is the causative agent of Coronavirus Disease 2019 (COVID-19), which has resulted in a devastating global pandemic with over 100 million cases and 2.4 million deaths worldwide (WHO, 2021). As SARS-CoV-2 has spread across the world, there has been a dramatic increase in the emergence of variants with mutations in the nonstructural and structural proteins. The viral spike protein is found on the outside of the virion and binds to the ACE2 receptor expressed on cells within the respiratory tract (Walls et al., 2020). As compared to the Wuhan-Hu-1 reference genome, several mutations within the spike protein have been identified over the past year. The first major spike protein variant to emerge was a mutation at position 614 from an Aspartic acid (D) to a Glycine (G). This mutation led to an increase in viral fitness, replication in the respiratory tract, binding to the ACE2 receptor, and confirmational changes within the spike protein (Gobeil et al., 2021; Ozono et al., 2021; Plante et al., 2020). Over the past few months, there has been a surge in the emergence of novel SARS-CoV-2 variants, raising significant concerns about alterations to viral fitness, transmission and disease. In particular, the emergence of the B.1.351 variant, which was originally identified in South Africa, includes several mutations within the structural and nonstructural proteins (Tegally et al., 2020).

Following SARS-CoV-2 infection in humans, antibody responses are rapidly generated against the viral spike protein (Suthar et al., 2020). The receptor binding motif within the spike protein interacts with the ACE2 receptor and is a major target of antibody-mediated neutralization. Longitudinal and cross-sectional studies have estimated that antibodies to the spike protein can last for at least a year following infection (Anand et al., 2021; Dan et al., 2021; Pradenas et al., 2021; Sherina et al., 2021). The mRNA-1273 vaccine encodes the viral spike protein and elicits a potent neutralizing antibody response to SARS-CoV-2 that is durable for several months (Anderson et al., 2020; Jackson et al., 2020; Widge et al., 2021). The emerging B.1.351 SARS-CoV-2 variant includes three mutations within the receptor-binding domain (K417N, E484K, N501Y) and several mutations within the spike protein which are likely to influence viral binding to the ACE2 receptor and resist neutralization by human immune sera (Greaney et al., 2021; Liu et al., 2021).

In this study, we compared antibody binding and viral neutralization against two variants that have emerged in various parts of the world. EHC-083E (herein referred to as the B.1 variant) is within the B.1 PANGO lineage and was isolated from a residual nasopharyngeal swab collected from a patient in Atlanta, GA in March 2020 (SARS-CoV-2/human/USA/GA-EHC-083E/2020). This variant contains the D614G mutation within the spike protein. The B.1.351 variant was isolated from an oropharyngeal swab from a patient in Ugu district, KwaZulu-Natal, South Africa in November 2020. The B.1.351 viral variant contains the following amino acid mutations within the viral spike protein: L18F, D80A, D215G, deletion at positions 242-244 (L242del, A243del, and L244del), K417N, E484K, N501Y and D614G. This virus was isolated as described by Sigal and colleagues (Wibmer et al., 2021). We subsequently plaque purified the virus followed by a single round of propagation in VeroE6 cells. Relative to the deposited sequence on GISAID (EPI_ISL_678615) we identified two additional mutations within the spike protein at positions Q677H and R682W (**Supplementary Figure 1**).

Following SARS-CoV-2 infection, antibody responses against the receptor binding domain (RBD) within the spike protein can be detected in most individuals around 8 days post-symptom onset (Suthar et al., 2020). Here, we analyzed a cohort of acutely infected COVID-19 patients (n=19) enrolled at Emory University Hospital, between 5-19 days after symptom onset (**Supplementary Table 1**). To determine if the RBD of the B.1.351 variant impacts IgG antibody binding, we utilized an electrochemiluminescence-based multiplex immune assay provided by Mesoscale Discovery (MSD). As compared to the B.1-lineage RBD-specific IgG responses (GMT: 4829; range: <239 – 168890), we found that all of the patients had significantly reduced IgG binding to the B.1.351 RBD (GMT: 1081; range: <239 – 20254). We next determined the impact on the neutralization capacity of these samples using a live virus neutralization assay. In comparison to the D614G variant (GMT: 135; range: <20 – 836), we observed a significant reduction in the neutralization capacity of samples from the acutely infected cohort against the B.1.351 variant (GMT: 40; range: <20 – 433). Of the samples that exhibited neutralization against the B.1 variant, we found that 4/15 samples (26%) failed to neutralize the B.1.351 variant. While there was a range of RBD-specific and neutralizing antibody responses across this cohort of acutely infected COVID-19 patients, we observed a stronger positive correlation of B.1-lineage RBD-specific IgG titers against the B.1 variant neutralization titers (R^2^= 0.47; *p*=0.0012; **Fig. 1C**) as compared to the B.1.351 RBD-specific IgG titers against the B.1.351 variant neutralization titers (R^2^= 0.27; *p*=0.02). This suggests that antibodies are capable of binding to the B.1.351 RBD, however, the mutations within the receptor-binding domain (K417N, E484K and N501Y) reduce the ability to neutralize the B.1.351 variant.

**Figure 1.**
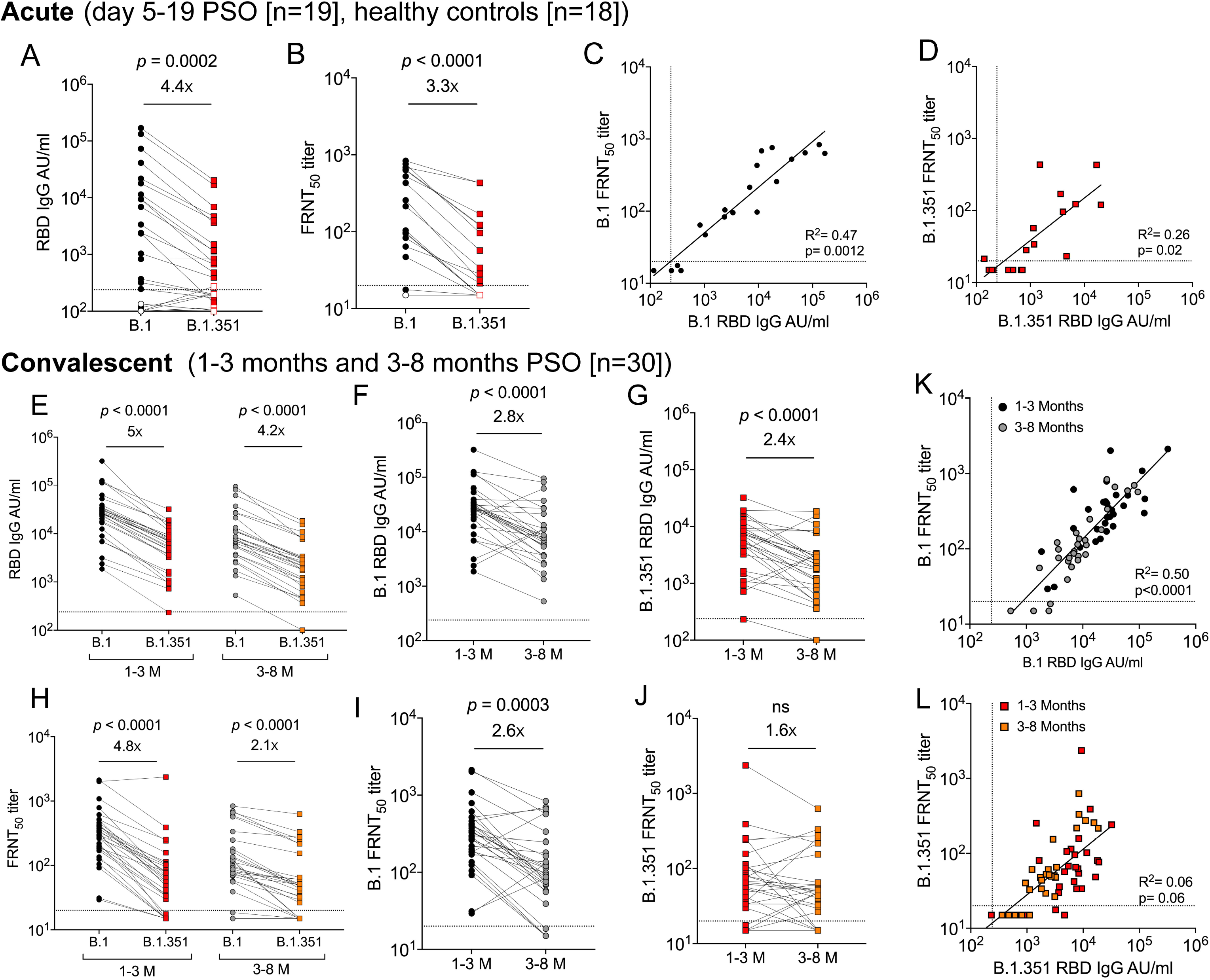
RBD binding and neutralizing antibody response against SARS-CoV-2 B.1.351 variant in SARS-CoV-2 infected individuals. Shown are data from the following cohorts based on natural infection: 19 acutely infected COVID-19 patients (5-19 days PSO; closed symbols), 30 convalescent COVID-19 individuals (1-3 months and 3-8 months PSO, closed symbols) and 18 healthy controls (open symbols). (A) IgG antibody responses against SARS-CoV-2 receptor binding domain (RBD) were measured by an electrochemiluminescent multiplex immunoassay and reported as arbitrary units per ml (AU/ml) as normalized by a standard curve for the B.1 (black) and B.1.351 (red) SARS-CoV-2 variants, (B) The 50% inhibitory titer (FRNT_50_) on the focus reduction neutralization test (FRNT) for the B.1 (black) and B.1.351 (red) SARS-CoV-2 variants, and correlations plots between the corresponding RBD and FRNT_50_ for the (C) B.1 variant and (D) B.1.351 variant are shown for the acutely infected COVID-19 patients. (E) Comparison of IgG antibody responses between the B.1 (black) and B.1.351 (red) SARS-CoV-2 variants at 1-3 month and the B.1 (grey) and B.1.351 (orange) SARS-CoV-2 variants at 3-8 month time points are shown for the convalescent COVID-19 patients. Changes in IgG antibody responses over two time points through 8 months for the (F) B.1 (1-3 months (black), 3-8 months (grey)) and (G) B.1.351 (1-3 months (red), 3-8 months (orange)) are shown for the convalescent COVID-19 patients. (H) Comparison of FRNT_50_ titer between the B.1 (black) and B.1.351 (red) SARS-CoV-2 variants at 1-3 month and the B.1 (grey) and B.1.351 (orange) SARS-CoV-2 variants at 3-8 month time points are shown for the convalescent COVID-19 patients. Changes in FRNT_50_ titers over two time points through 8 months for the (I) B.1 (1-3 months (black), 3-8 months (grey)) and (J) B.1.351 (1-3 months (red), 3-8 months (orange)) are shown for the convalescent COVID-19 patients. Correlations plots between the corresponding RBD and FRNT_50_ for the (K) B.1 variant (1-3 month (black), 3-8 month (grey)) and (L) B.1.351 variant (1-3 month (red), 3-8 month (orange)) are shown for the convalescent COVID-19 patients. The dotted line in the RBD binding assays represents the limit of detection (239 IgG AU/ml). The dotted line in the FRNT assays represents the maximum concentrations of the serum tested (1/20). Statistical significance was determined using a Wilcoxon paired t-test. The GMT fold change for the respective isolates relative to B.1 is shown in each of the plots. Correlation analysis was performed by log transformation of the RBD-specific IgG AU/ml values or FRNT_50_ titers followed by linear regression analysis.

Recent studies have found that binding and neutralizing antibodies are maintained for at least eight months following SARS-CoV-2 infection (Dan et al., 2021; Pradenas et al., 2021; Sherina et al., 2021). To understand how antibody breadth is impacted during convalescence, we performed a longitudinal analysis of RBD binding and viral neutralization in 30 convalescent COVID-19 individuals across two longitudinally sampled timepoints through 8 months (**Supplementary Table 2**). We observed a significant reduction in IgG binding to the B.1.351 RBD at the 1-3 month timepoint (B.1: GMT: 24000; range: 1856 – 320059; B.1.351: GMT: 4792; range: <239 – 32158) and 3-8 month timepoint (B.1: GMT: 8314; range: 527 – 94643; B.1.351: GMT: 1946; range: <239 – 18544; **Figure 1E**). We observed similar reductions in IgG binding titers to the B.1 and B.1.351 RBD across these two timepoints (**Figure 1F-G**). We next determined the impact on the neutralization capacity of these samples across the two timepoints. At the 1–3-month timepoint, we observed a 4.8-fold reduction (*p*<0.0001) in neutralization capacity between the B.1 variant (GMT: 288; range: 29 – 2117) and the B.1.351 variant (GMT: 59; range: <20 – 2363). At the 3–8-month timepoint, we observed a 2.1-fold reduction (*p*<0.0001) in neutralization capacity between the B.1 variant (GMT: 107; range: <20 – 836) and the B.1.351 variant (GMT: 50; range: <20 – 627). Of the samples that exhibited neutralization against the D614G variant, 7 of 30 samples (23%) at the 1–3-month timepoint and 4 of 26 samples (15%) at the 3–8-month timepoint failed to neutralize the B.1.351 variant. Regression analysis showed a significant correlation between B.1-lineage RBD-specific IgG and neutralization titers against the B.1 variant (R^2^= 0.5; *p*<0.0001; **Fig 1K**). Unlike the acutely infected COVID-19 patients, the convalescent COVID-19 infected individuals did not show a correlation between the B.1.351 RBD-specific IgG and neutralization titers against the B.1.351 variant (R^2^= 0.06; *p*=0.06; **Fig 1L**). Taken together, these data demonstrate that antibody titers are reduced through 8 months following SARS-CoV-2 infection, however, there is a modest impact on the neutralization potency against the B.1.351 variant.

The messenger RNA vaccine, mRNA-1273, generates durable neutralizing antibodies against SARS-CoV-2 (Anderson et al., 2020). We examined binding and neutralizing antibody titers in 19 healthy adult participants that received two injections of the mRNA-1273 vaccine at a dose of 100 μg (age >56 years; 14 days post-2nd dose; **Supplementary Table 2**). We found that all vaccinated individuals had significantly reduced IgG binding to the B.1.351 RBD (GMT: 83909; range: 2588 – 333451) as compared to the B.1-lineage RBD-specific IgG responses (GMT: 316554; range: 7313 – 975553; **Fig 2A**). Similarly, we observed a 3.8-fold reduction (*p*<0.0001) in neutralization capacity between the B.1. variant (GMT: 734; range: 256 – 2868) and the B.1.351 variant (GMT: 191; range: 61 – 830; **Fig 2B**). In contrast to the infected individuals, all vaccinated individuals retained neutralization capacity against the B.1.351 variant. Further, we observed a strong correlation between the corresponding RBD-specific IgG titers to the B.1 variant neutralization titers (R^2^= 0.75; *p*<0.0001; **Fig 2C**) and the B.1.351 variant neutralization titers (R^2^= 0.85; *p*<0.0001). These findings demonstrate that the antibodies elicited by the mRNA-1273 vaccine are effective at neutralizing the B.1.351 variant.

**Figure 2.**
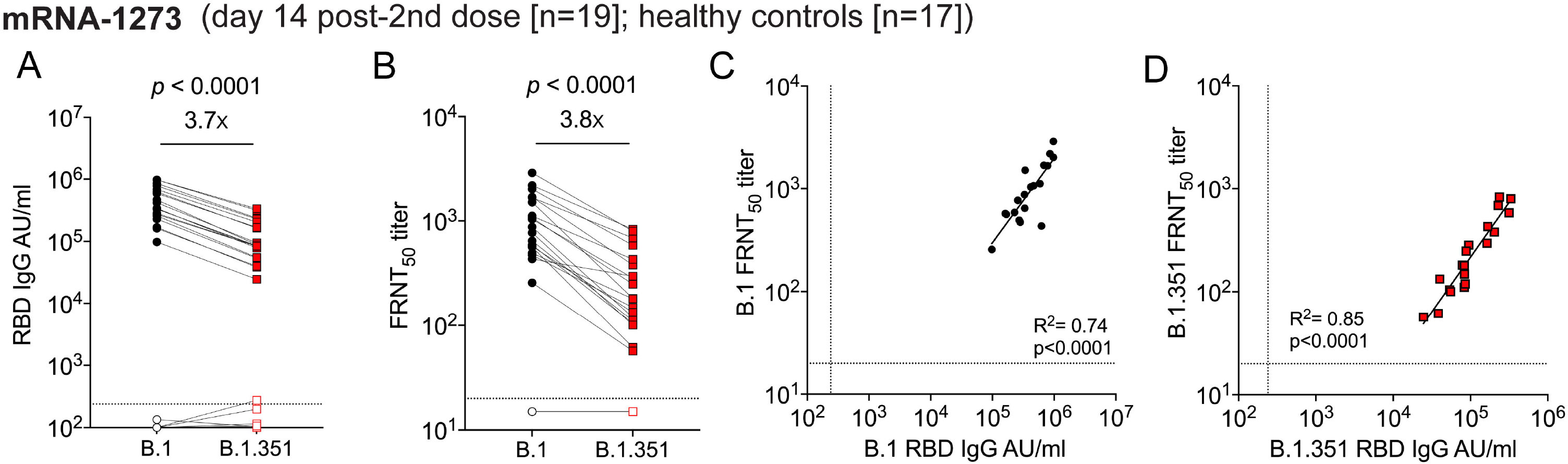
RBD binding and neutralizing antibody response against SARS-CoV-2 B.1.351 viral variant among mRNA-1273 vaccinated individuals. Shown are data from the individuals that received 100 μg of mRNA-1273 on day 14 post-2nd dose (>56 years or older, 19 participants; closed symbols) and 18 healthy controls (open symbols). (A) IgG antibody responses against SARS-CoV-2 receptor binding domain (RBD) were measured by an electrochemiluminescent multiplex immunoassay and reported as arbitrary units per ml (AU/ml) as normalized by a standard curve, for the B.1 (black) and B.1.351 (red) SARS-CoV-2 variants (B) The 50% inhibitory titer (FRNT_50_) on the focus reduction neutralization test (FRNT) for the B.1 (black) and B.1.351 (red) SARS-CoV-2 variants, and correlations plots between the corresponding RBD and FRNT_50_ for the (C) B.1 variant and (D) B.1.351 variant are shown. The dotted line in the RBD binding assays represents the limit of detection (239 AU/ml). The dotted line in the FRNT assays represents the maximum concentrations of the serum tested (1/20). Statistical significance was determined using a Wilcoxon paired t-test. The GMT fold change for the respective isolates relative to B.1 is shown in each of the plots. Correlation analysis was performed by log transformation of the RBD-specific IgG AU/ml values or FRNT_50_ titers followed by linear regression analysis.

This study examined the impact of infection- and vaccine-induced antibody responses against two SARS-CoV-2 variants. We observed reduced antibody binding to the B.1.351-derived RBD of the spike protein and neutralization potency against the B.1.351 variant virus in sera from SARS-CoV-2 infected and vaccinated individuals. Using our longitudinal convalescent COVID-19 cohort, we examined the impact on antibody binding to the RBD and viral neutralization across the SARS-CoV-2 variants. One of the interesting findings is that in most convalescent COVID-19 individuals, we observed less of an impact on viral neutralization against the B.1.351 variant at longer periods after infection. This suggests that antibodies capable of neutralizing the B.1.351 variant are generated early during infection and are durable for several months.

The immune correlates of protection against SARS-CoV-2 are not yet known. We and others have previously shown, IgG antibody responses to the RBD can serve as a surrogate of viral neutralization in infected individuals (Greaney et al., 2021; Piccoli et al., 2020; Suthar et al., 2020). However, the B.1.351 contains three mutations (K417N, E484K and N501Y) within the RBD, which, combined, likely impact antibody binding and viral neutralization. Of these mutations, we and others have shown that the presence of N501Y mutation within the RBD in B.1.1.7 UK variant does not affect the neutralizing ability of serum from either naturally infected or mRNA-1273 vaccinated individuals (Edara et al., 2021; Johnson et al., 2021; Rathnasinghe et al., 2021; Shen et al., 2021; Wu et al., 2021). The substitution at position E484, located in the receptor-binding ridge epitope (Greaney et al., 2021), shows resistance to the neutralization of convalescent human sera. Single point mutant pseudoviruses, chimeric viruses, or recombinant infectious clone-derived SARS-CoV-2 have demonstrated that this mutation displays resistance to neutralization by infection- and vaccine-induced antibodies (Johnson et al., 2021; Liu et al., 2021; Shen et al., 2021; Xie et al., 2021). This suggests that a majority of individuals develop antibodies that target this region within the RBD. However, it is still unclear whether this mutation also impacts viral fitness, pathogenesis or transmission.

We observed that most of the sera samples from acute and convalescent COVID-19 individuals showed antibody binding to the B.1.351-dervied RBD. In addition to most of these samples showing capacity to neutralize the B.1.351 variant, the effector functions of these antibodies could also contribute to controlling SARS-CoV-2 infection. Recent studies have shown that antibody Fc effector functions are important for mediating protection against SARS-CoV-2 in mouse and hamster models (Schäfer et al., 2021; Winkler et al., 2020). Future studies should evaluate the contribution of Fc effector functions in promoting viral control and protective immunity following infection or vaccination against SARS-CoV-2.

One of the limitations is that our study focused on antibody binding to the RBD of the spike protein. It is becoming increasingly clear that monoclonal antibodies targeting the N-terminal domain and other regions of the spike protein outside of the RBD can neutralize SARS-CoV-2 (Greaney et al., 2021; Liu et al., 2021; Suryadevara et al., 2021). Examining binding to the full-length B.1.351 spike protein, as well as individual point mutations, will provide important insight to the breadth of the antibody response to the viral spike protein following virus infection and vaccination. Another limitation is that the B.1.351 virus that we used in our study contains two substitutions within the spike protein that were not reported in the reference sequence deposited into GISAID (EPI_ISL_678615). One of these is a substitution of a Q677H, which has now been reported in multiple lineages of circulating variants of SARS-CoV-2 in the US population as early as mid-August 2020 (Hodcroft et al., 2021). The other is a substitution (R682W) within the furin cleavage motif (PRRAR) located between the S1/S2 regions of the spike protein.

Our results show that despite few fold decrease, most infected individuals showed binding and neutralizing titers against the B.1.351 variant in acute and convalescent sera, and further, all mRNA-1273 vaccinated individuals still maintained neutralization. These findings support the notion that in the context of the B.1.351 variant, infection- and vaccine-induced immunity can provide protection against COVID-19.

## Acknowledgments

This work was supported in part by grants (P51 OD011132, 3U19AI057266-17S1, U19AI090023, R01AI127799, R01AI148378, K99AI153736, 1UM1AI148576-01, 5R38AI140299-03 and UM1AI148684 to Emory University) from the National Institute of Allergy and Infectious Diseases (NIAID), National Institutes of Health (NIH), by the Emory Executive Vice President for Health Affairs Synergy Fund award, the Pediatric Research Alliance Center for Childhood Infections and Vaccines and Children’s Healthcare of Atlanta, COVID-Catalyst-I^3^ Funds from the Woodruff Health Sciences Center and Emory School of Medicine, Woodruff Health Sciences Center 2020 COVID-19 CURE Award, and the Vital Projects/Proteus funds. We also thank Jim Wilbur (Mesoscale Discovery) for providing reagents to perform the RBD-binding assays. The following reagent was obtained through BEI Resources, NIAID, NIH: SARS-Related Coronavirus 2, Isolate hCoV-19/South Africa/KRISP-K005325/2020, NR-54009, contributed by Alex Sigal and Tulio de Oliveira. We thank Natalie Thornburg, Clinton Paden, and Suxiang Tong for sequencing and analysis of the B.1.351 variant (CDC, Atlanta, GA).

---

**Supplemental Figure 1.** (A) Structure of SARS-CoV-2 spike protein (single monomer is shown) (Walls et al., 2020) with the mutations highlighted in red. Additional mutations Q677H and R682W that are not reported in the GISAID reference sequence (EPI_ISL_678615) were highlighted in green. (B) A schematic of the amino acid changes within the spike protein are shown between the SARS-CoV-2 variants.

## STAR METHODS

KEY RESOURCES TABLE.

| REAGENT or RESOURCE | SOURCE | IDENTIFIER |
| --- | --- | --- |
| **Antibodies** |  |  |
| CR3022-biotin | Dr. Jens Wrammert<br>Emory University |  |
| **Virus Strains** |  |  |
| SARS-CoV-2/human/USA/GA-EHC-083E/2020 (EHC-083E) | residual nasopharyngeal swab |  |
| N501Y HV2 (B.1.351) | BEI Resources |  |
| **Biological Samples** |  |  |
| Acute and Convalescent human Serum/Plasma samples | Emory University Hospital |  |
| mRNA-1273 Phase-1 study samples | Division of Microbiology and Infectious Diseases, NIAID |  |
| **Chemicals, Peptides, and Recombinant Proteins** |  |  |
| Methylcellulose | Sigma-Aldrich | Cat. #: M0512-250G |
| TrueBlue Peroxidase Substrate | KPL | Cat. #: 5510-0050 |
| **Experimental Models: Cell Lines** |  |  |
| VeroE6 C1008 cells | ATCC | Cat# CRL-1586, RRID:CVCL_0574 |
| **Software and Algorithms** |  |  |
| GraphPad Prism (v7 and v8) | N/A | N/A |

### RESOURCE AVAILABILITY

#### Lead Contact

Further information and requests for resources and reagents should be directed to and will be fulfilled by the Lead Contact Author Mehul Suthar.

#### Materials availability

All unique/stable reagents generated in this study are available from the Lead Contact with a completed Materials Transfer Agreement.

#### Data and code availability

The datasets supporting the current study are available from the corresponding author on request.

### EXPERIMENT MODEL AND SUBJECT DETAILS

#### Ethics Statement

For samples Emory University, collection and processing were performed under approval from the University Institutional Review Board (IRB #00001080 and #00022371). Adults ≥18 years were enrolled who met eligibility criteria for SARS-CoV-2 infection (PCR confirmed by a commercially available assay) and provided informed consent. For the mRNA-1273 phase 1 clinical trial, the neutralization assays were conducted on deidentified specimens, as protocol-defined research. The mRNA-1273 phase 1 clinical trial (NCT04283461) was reviewed and approved by the Advarra institutional review board, which functioned as a single board. The trial was overseen by an independent safety monitoring committee. All participants provided written informed consent before enrollment. The trial was conducted under an Investigational New Drug application submitted to the Food and Drug Administration. The NIAID served as the trial sponsor and made all decisions regarding the study design and implementation.

#### Serum samples

For Emory University, acute peripheral blood samples were collected from hospitalized patients at the time of enrollment. Convalescent samples from COVID-19 patients were collected when the patients were able to return for a visit to the clinical research site at the next study visit. Convalescent samples were collected at a range of times (1-8 months) post symptom onset. Serum samples for the mRNA-1273 phase 1 study were obtained from the Division of Microbiology and Infectious Diseases, National Institute of Allergy and Infectious diseases for the mRNA-1273 phase 1 study team and Moderna Inc. Study protocols and results were previously reported (Anderson et al., 2020). Samples tested were collected from 19 healthy individuals on day 14 post-2^nd^ dose of the mRNA-1273 vaccine.

#### Cells

VeroE6 cells were obtained from ATCC (clone E6, ATCC, #CRL-1586) and cultured in complete DMEM medium consisting of 1x DMEM (VWR, #45000-304), 10% FBS, 25mM HEPES Buffer (Corning Cellgro), 2mM L-glutamine, 1mM sodium pyruvate, 1x Non-essential Amino Acids, and 1x antibiotics.

#### Virus isolation and sequencing

EHC-083E (herein referred to as the B.1 variant) was derived from a residual nasopharyngeal swab collected from an Emory Healthcare patient in March 2020, as part of a study approved by the institutional review board at Emory University. As described previously (Babiker et al., 2020), the primary sample underwent RNA extraction, DNAse treatment, random primer cDNA synthesis, Nextera XT tagmentation, Illumina sequencing, and reference-based viral genome assembly. Results were confirmed by sequencing of an independent library. A total of 47,542,787 reads were derived from this sample, leading to 100% SARS-CoV-2 genome coverage with a mean depth of 488X. All sequencing reads (cleaned of human reads) are available on NCBI under BioProject PRJNA634356, and the consensus SARS-CoV-2 genome is available under GenBank accession number MW008579.1. Following virus isolation, culture supernatant underwent metagenomic sequencing as described above. A total of 836,424 paired-end 150bp reads were generated by Illumina MiSeq, and reference-based SARS-CoV-2 genome assembly was performed using viral-ngs v.2.1.7 (https://github.com/broadinstitute/viral-ngs) with reference sequence NC_045512.2. The resulting consensus sequence had 100% coverage with a mean depth of 750X and was identical to the consensus sequence from the primary sample. The B.1 variant was plaque purified on VeroE6 cells propagated two times on VeroE6 cells (MOI 0.01), aliquoted to generate a working stock and sequenced. The B.1.351 variant was isolated as previously described (Tegally et al., 2020). Our laboratory plaque isolated the virus on VeroE6 cells followed by a single round of propagation on VeroE6 cells (MOI 0.05), aliquoted to generate a working stock and sequenced. As described (https://github.com/CDCgov/SARS-CoV-2_Sequencing/blob/master/protocols/CDC-Comprehensive/CDC_SARS-CoV-2_Sequencing_200325-2.pdf) the primary sample underwent RNA extraction and cDNA synthesis was performed with random primers followed by pooling non-overlapping amplicons and Barcoding and library prep with ONT Ligation protocol and 96 PCR Barcoding expansion. Quality check was performed by excluding reads that are not in 200-800 base range. The resulting sequences were mapped to Wuhan reference with minimap2. Soft clip primer regions were identified using BAMClipper based on mapping position. Consensus variants were identified using ONT Medaka software and the variants were filtered with < 30 g score. Finally, variants and masking were applied to the reference sequence. Viral titers were determined by focus-forming assay on VeroE6 cells. Viral stocks were stored at −80°C until use.

#### RBD-binding assay

Plasma from acute and convalescent COVID-19 patients, mRNA-1273 vaccine recipients (14 days post-dose 2), and healthy controls was tested for IgG antibody binding against SARS-CoV-2 reference RBD (herein referred to as the B.1-lineage RBD), and B.1.351 RBD using an electrochemiluminescent-based multiplex immunoassay (kindly provided by Mesoscale Discovery (MSD)). Plates pre-coated with the RBD antigens were blocked for 30 minutes at room temperature, shaking at a speed of 700 rpm, with 150 uL per well of MSD Blocker A. To assess IgG binding, plasma samples were diluted 1:5000 and MSD Reference Standard-1 was diluted per MSD instructions in MSD Diluent 100. 50 uL of each sample and Reference Standard-1 dilution was added to the plates and incubated for two hours at room temperature, shaking at a speed of 700 rpm. Following this, 50 uL per well of 1X MSD SULFO-TAG™ Anti-Human IgG Antibody was added and incubated for one hour at room temperature, shaking at a speed of 700 rpm. Following the detection reagent step, 150 uL per well of MSD Gold™ Read Buffer B was added to each plate immediately prior to reading on an MSD plate reader. Plates were washed three times with 300 uL PBS/0.05% Tween between each step. Data was analyzed using Discovery Workbench and GraphPad Prism software. Plasma antibody concentration in arbitrary units (AU) was calculated relative to Reference Standard 1.

#### Focus Reduction Neutralization Assay

FRNT assays were performed as previously described (Vanderheiden et al., 2020). Briefly, samples were diluted at 3-fold in 8 serial dilutions using DMEM (VWR, #45000-304) in duplicates with an initial dilution of 1:10 in a total volume of 60 μl. Serially diluted samples were incubated with an equal volume of SARS-CoV-2 (100-200 foci per well) at 37° C for 1 hour in a round-bottomed 96-well culture plate. The antibody-virus mixture was then added to Vero cells and incubated at 37° C for 1 hour. Post-incubation, the antibody-virus mixture was removed and 100 μl of prewarmed 0.85% methylcellulose (Sigma-Aldrich, #M0512-250G) overlay was added to each well. Plates were incubated at 37° C for 24 hours. After 24 hours, methylcellulose overlay was removed, and cells were washed three times with PBS. Cells were then fixed with 2% paraformaldehyde in PBS (Electron Microscopy Sciences) for 30 minutes. Following fixation, plates were washed twice with PBS and 100 μl of permeabilization buffer (0.1% BSA [VWR, #0332], Saponin [Sigma, 47036-250G-F] in PBS), was added to the fixed Vero cells for 20 minutes. Cells were incubated with an anti-SARS-CoV spike primary antibody directly conjugated to biotin (CR3022-biotin) for 1 hour at room temperature. Next, the cells were washed three times in PBS and avidin-HRP was added for 1 hour at room temperature followed by three washes in PBS. Foci were visualized using TrueBlue HRP substrate (KPL, # 5510-0050) and imaged on an ELISPOT reader (CTL).

#### Quantification and Statistical Analysis

Antibody neutralization was quantified by counting the number of foci for each sample using the Viridot program (Katzelnick et al., 2018). The neutralization titers were calculated as follows: 1 - (ratio of the mean number of foci in the presence of sera and foci at the highest dilution of respective sera sample). Each specimen was tested in duplicate. The FRNT-50 titers were interpolated using a 4-parameter nonlinear regression in GraphPad Prism 8.4.3. Samples that do not neutralize at the limit of detection at 50% are plotted at 15 and was used for geometric mean calculations. The SARS-CoV-2 Spike structure was visualized with PyMOL (Schrödinger, Inc.).

