## Supplementary figures and images for "Reduced binding and neutralization of infection- and vaccine-induced antibodies to the B.1.351 (South African) SARS-CoV-2 variant"

### Supplementary Figure 1

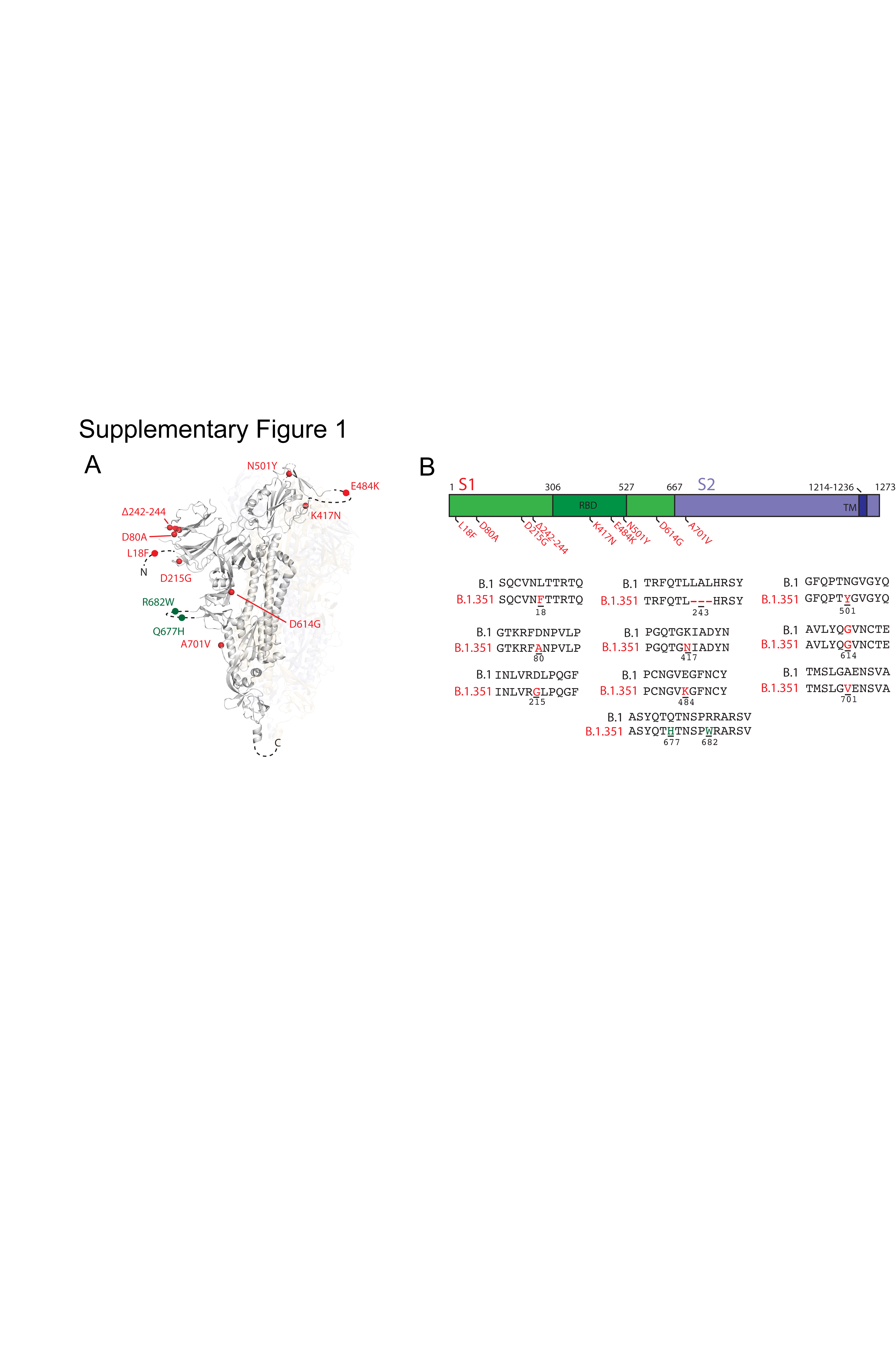
