## Supplementary Table 1 for "Reduced binding and neutralization of infection- and vaccine-induced antibodies to the B.1.351 (South African) SARS-CoV-2 variant"

**Table 1. COVID-19 patient cohort and healthy controls**

| **GROUP** | **ID#** | **Date samples collected** | **Days After Symptom Onset** | **B.1 RBD IgG Binding** | **B.1 FRNT50** | **B.1.351 RBD IgG Binding** | **B.1.351 FRNT50** |
| --- | --- | --- | --- | --- | --- | --- | --- |
| ACUTE | 1 | 7/7/2020 | 5 | 246 | <20 | 170 | <20 |
|  | 2 | 7/7/2020 | 12 | 830 | 64 | 838 | 28 |
|  | 3 | 7/7/2020 | 11 | 1038 | 47 | 391 | <20 |
|  | 4 | 7/8/2020 | 7 | 2352 | 83 | 140 | 21 |
|  | 5 | 7/9/2020 | 11 | 2397 | 104 | 671 | <20 |
|  | 6 | 7/10/2020 | 11 | 9425 | 97 | 709 | <20 |
|  | 7 | 7/10/2020 | 9 | 9380 | 431 | 1136 | 57 |
|  | 8 | 7/10/2020 | 10 | 41146 | 526 | 3625 | 170 |
|  | 9 | 7/14/2020 | 7 | 132731 | 836 | 16795 | 430 |
|  | 10 | 7/14/2020 | 11 | 3370 | 95 | 477 | <20 |
|  | 11 | 7/15/2020 | 11 | 6866 | 214 | 1169 | 34 |
|  | 12 | 7/17/2020 | 11 | 11395 | 685 | 1499 | 433 |
|  | 13 | 7/21/2020 | 10 | 21654 | 257 | 4672 | 23 |
|  | 14 | 7/23/2020 | 15 | 168890 | 633 | 20254 | 120 |
|  | 15 | 7/23/2020 | 8 | 369 | <20 | 199 | <20 |
|  | 16 | 7/23/2020 | 9 | 116 | <20 | 205 | <20 |
|  | 17 | 7/28/2020 | 14 | 72783 | 642 | 6875 | 122 |
|  | 18 | 7/28/2020 | 19 | 318 | 18 | 193 | <20 |
|  | 19 | 7/28/2020 | 14 | 17903 | 760 | 4026 | 96 |
| Convalescent | 20 | 4/17/2020 | 34 | 20552 | 136 | 4669 | <20 |
|  |  | 10/14/2020 | 214 | 2668 | 19 | 467 | <20 |
|  | 21 | 4/22/2020 | 44 | 17974 | 182 | 3767 | 36 |
|  |  | 10/16/2020 | 221 | 61921 | 592 | 15603 | 257 |
|  | 22 | 4/22/2020 | 31 | 34856 | 288 | 8353 | 54 |
|  |  | 10/2/2020 | 194 | 6319 | 58 | 2157 | 29 |
|  | 23 | 4/22/2020 | 37 | 3126 | 31 | 990 | <20 |
|  |  | 10/29/2020 | 227 | 11202 | 84 | 3130 | 26 |
|  | 24 | 4/23/2020 | 39 | 24861 | 218 | 3230 | 18 |
|  |  | 10/5/2020 | 204 | 21307 | 179 | 1732 | 48 |
|  | 25 | 4/23/2020 | 40 | 112991 | 1086 | 32158 | 241 |
|  |  | 10/28/2020 | 228 | 27281 | 337 | 11032 | 274 |
|  | 26 | 4/27/2020 | 30 | 1856 | 92 | 232 | <20 |
|  |  | 10/27/2020 | 213 | 527 | <20 | 83 | <20 |
|  | 27 | 4/28/2020 | 46 | 29979 | 266 | 9409 | 34 |
|  |  | 10/2/2020 | 203 | 5536 | 68 | 1810 | 34 |
|  | 28 | 4/28/2020 | 42 | 6878 | 614 | 1466 | 253 |
|  |  | 10/2/2020 | 199 | 1707 | 56 | 442 | <20 |
|  | 29 | 4/29/2020 | 50 | 26483 | 789 | 5446 | 68 |
|  |  | 11/18/2020 | 253 | 5752 | 76 | 1228 | 61 |
|  | 30 | 4/30/2020 | 48 | 63027 | 513 | 16597 | 48 |
|  |  | 10/21/2020 | 222 | 8336 | 80 | 2391 | 61 |
|  | 31 | 5/6/2020 | 47 | 12499 | 335 | 1083 | <20 |
|  |  | 10/16/2020 | 210 | 8106 | 114 | 809 | <20 |
|  | 32 | 5/8/2020 | 56 | 27035 | 263 | 7122 | 96 |
|  |  | 10/28/2020 | 229 | 11531 | 112 | 3370 | 65 |
|  | 33 | 5/13/2020 | 61 | 28451 | 348 | 8116 | 62 |
|  |  | 10/5/2020 | 206 | 8214 | 71 | 3048 | 50 |
|  | 34 | 5/15/2020 | 60 | 8923 | 105 | 934 | <20 |
|  |  | 10/15/2020 | 213 | 2511 | <20 | 360 | <20 |
|  | 35 | 5/21/2020 | 57 | 6844 | 187 | 1642 | 80 |
|  |  | 10/15/2020 | 204 | 3702 | 98 | 938 | 40 |
|  | 36 | 7/29/2020 | 31 | 2379 | 29 | 720 | <20 |
|  |  | 12/15/2020 | 117 | 11025 | 130 | 3222 | 48 |
|  | 37 | 8/21/2020 | 52 | 23318 | 417 | 4675 | 57 |
|  |  | 2/2/2021 | 217 | 3728 | 76 | 1122 | 33 |
|  | 38 | 4/29/2020 | 57 | 34517 | 199 | 6880 | 42 |
|  |  | 9/18/2020 | 199 | 5338 | 39 | 1115 | <20 |
|  | 39 | 5/18/2020 | 45 | 16915 | 125 | 3671 | 30 |
|  |  | 10/1/2020 | 181 | 1345 | <20 | 472 | <20 |
|  | 40 | 6/15/2020 | 68 | 25376 | 423 | 7383 | 65 |
|  |  | 10/15/2020 | 190 | 7051 | 92 | 2197 | 53 |
|  | 41 | 5/11/2020 | 68 | 320059 | 2117 | 13385 | 390 |
|  |  | 9/28/2020 | 208 | 94643 | 570 | 7697 | 218 |
|  | 42 | 6/19/2020 | 48 | 53032 | 373 | 17924 | 80 |
|  |  | 10/13/2020 | 164 | 6901 | 87 | 2712 | 49 |
|  | 43 | 8/14/2020 | 83 | 30934 | 2014 | 9345 | 2363 |
|  |  | 11/20/2020 | 181 | 26653 | 837 | 8386 | 627 |
|  | 44 | 8/21/2020 | 50 | 32703 | 328 | 6095 | 114 |
|  |  | 10/2/2020 | 92 | 3528 | 121 | 614 | <20 |
|  | 45 | 8/21/2020 | 38 | 125717 | 460 | 19116 | 77 |
|  |  | 12/16/2020 | 155 | 81491 | 697 | 18544 | 216 |
|  | 46 | 8/21/2020 | 76 | 26864 | 399 | 5127 | 106 |
|  |  | 12/15/2020 | 192 | 7648 | 162 | 1824 | 44 |
|  | 47 | 8/26/2020 | 58 | 26400 | 170 | 7650 | 34 |
|  |  | 1/6/2021 | 191 | 8647 | 136 | 2477 | 52 |
|  | 48 | 8/26/2020 | 42 | 122719 | 298 | 11616 | 102 |
|  |  | 1/12/2021 | 191 | 13285 | 248 | 2914 | 154 |
|  | 49 | 10/9/2020 | 91 | 38928 | 518 | 8354 | 157 |
|  |  | 1/6/2021 | 180 | 37097 | 664 | 8431 | 331 |
| Healthy Controls | 50 | 8/19/2020 | NA | 100 | <20 | 100 | <20 |
|  | 51 | 8/19/2020 | NA | 605 | <20 | 477 | <20 |
|  | 52 | 8/18/2020 | NA | 100 | <20 | 100 | <20 |
|  | 53 | 8/18/2020 | NA | 100 | <20 | 100 | <20 |
|  | 54 | 8/18/2020 | NA | 100 | <20 | 100 | <20 |
|  | 55 | 8/18/2020 | NA | 100 | <20 | 100 | <20 |
|  | 56 | 8/18/2020 | NA | 100 | <20 | 100 | <20 |
|  | 57 | 8/18/2020 | NA | 100 | <20 | 100 | <20 |
|  | 58 | 8/18/2020 | NA | 100 | <20 | 198 | <20 |
|  | 59 | 8/18/2020 | NA | 100 | <20 | 100 | <20 |
|  | 60 | 8/18/2020 | NA | 100 | <20 | 100 | <20 |
|  | 61 | 8/18/2020 | NA | 100 | <20 | 275 | <20 |
|  | 62 | 8/18/2020 | NA | 100 | <20 | 100 | <20 |
|  | 63 | 8/18/2020 | NA | 100 | <20 | 106 | <20 |
|  | 64 | 6/1/2015 | NA | 100 | <20 | 115 | <20 |
|  | 65 | 1/31/2017 | NA | 134 | <20 | 104 | <20 |
|  | 66 | 2/1/2017 | NA | 100 | <20 | 100 | <20 |
|  | 67 | 10/23/2012 | NA | 100 | <20 | 100 | <20 |
