## Supplementary Table 2 for "Reduced binding and neutralization of infection- and vaccine-induced antibodies to the B.1.351 (South African) SARS-CoV-2 variant"

**Table 2. mRNA-1273 vaccine cohort**

| **ID#** | **B.1 RBD IgG Binding** | **B.1 FRNT50** | **B.1.351 RBD IgG Binding** | **B.1.351 FRNT50** |
| --- | --- | --- | --- | --- |
| 1 | 338224 | 1507 | 78391 | 182 |
| 2 | 282384 | 472 | 83731 | 149 |
| 3 | 855722 | 2176 | 237047 | 831 |
| 4 | 419023 | 1037 | 94860 | 284 |
| 5 | 334576 | 645 | 83292 | 111 |
| 6 | 228290 | 589 | 53645 | 105 |
| 7 | 975553 | 2009 | 315578 | 584 |
| 8 | 330250 | 876 | 85225 | 119 |
| 9 | 779702 | 1669 | 228795 | 686 |
| 10 | 168570 | 562 | 38354 | 62 |
| 11 | 623291 | 434 | 163678 | 297 |
| 12 | 971861 | 2868 | 333451 | 797 |
| 13 | 464656 | 1066 | 88003 | 249 |
| 14 | 682520 | 1685 | 203796 | 379 |
| 15 | 271291 | 493 | 83435 | 180 |
| 16 | 97844 | 257 | 24634 | 57 |
| 17 | 161831 | 573 | 39988 | 133 |
| 18 | 258182 | 770 | 55284 | 100 |
| 19 | 591530 | 1115 | 167095 | 429 |
